# Comparison of Transgenic and Adenovirus hACE2 Mouse Models for SARS-CoV-2 Infection

**DOI:** 10.1101/2020.07.06.190066

**Authors:** Raveen Rathnasinghe, Shirin Strohmeier, Fatima Amanat, Virginia L. Gillespie, Florian Krammer, Adolfo García-Sastre, Lynda Coughlan, Michael Schotsaert, Melissa Uccellini

## Abstract

Severe acute respiratory syndrome CoV-2 (SARS-CoV-2) is currently causing a worldwide pandemic with high morbidity and mortality. Development of animal models that recapitulate important aspects of coronavirus disease 2019 (COVID-19) is critical for the evaluation of vaccines and antivirals, and understanding disease pathogenesis. SARS-CoV-2 has been shown to use the same entry receptor as SARS-CoV-1, human angiotensin-converting enzyme 2 (hACE2)(1-3). Due to amino acid differences between murine and hACE2, inbred mouse strains fail to support high titer viral replication of SARS-CoV-2 virus. Therefore, a number of transgenic and knock-in mouse models, as well as viral vector-mediated hACE2 delivery systems have been developed. Here we compared the K18-hACE2 transgenic model to adenovirus-mediated delivery of hACE2 to the mouse lung. We show that K18-hACE2 mice replicate virus to high titers in both the lung and brain leading to lethality. In contrast, adenovirus-mediated delivery results in viral replication to lower titers limited to the lung, and no clinical signs of infection with a challenge dose of 10^4^ plaque forming units. The K18-hACE2 model provides a stringent model for testing the ability of vaccines and antivirals to protect against disease, whereas the adenovirus delivery system has the flexibility to be used across multiple genetic backgrounds and modified mouse strains.

## Main text introduction

A novel coronavirus, severe acute respiratory syndrome coronavirus 2 (SARS-CoV-2) emerged in China in December 2019 (4), and has rapidly spread throughout the world. The virus likely originated in bats (5, 6), before potentially jumping to an intermediate host and then to humans. From December 2019 to July 2020, SARS-CoV-2 has caused over 10 million infections and 500,000 deaths worldwide. The virus can cause asymptomatic, mild, or severe respiratory infection in different individuals – and can transmit from asymptomatic or presymptomatic individuals, making containment with public health measures very difficult. The disease caused by the virus has been named coronavirus disease 2019 (COVID-19). Mild cases of COVID-19 are characterized by fever, cough, and fatigue, while severe cases involve bilateral interstitial pneumonia, and cardiac and clotting complications (7-10). No specific vaccines are currently available (11). Recently, the antiviral remdesivir has been approved for the treatment of COVID-19 patients, but its efficacy and supply is limited (12, 13). However, an unprecedented response from the research community has resulted in numerous therapeutics and vaccine candidates being investigated both pre-clinically (14-18) and clinically, with several clinical trials now underway (19, 20).

SARS-CoV-2 is highly similar to SARS-CoV-1, which also emerged via interspecies transmission in 2003 (21). Both viruses bind to host cells via the interaction of the viral spike (S) protein with human angiotensin-converting enzyme 2 (hACE2) (2, 3, 22-24). A number of amino acid changes in SARS-CoV-2 S increase the affinity of binding to hACE2 (25). Species-specific differences in ACE2 are a critical determinant for S protein binding and high-titer viral replication in animal models. The mouse ortholog of ACE2 is incompatible with S, therefore typical inbred mouse strains do not support viral replication. This is a major challenge, as a large amount of pre-clinical vaccine, anti-viral, and therapeutic monoclonal antibody (mAb) testing is usually initially performed in mice, due to the availability of reagents for immunology studies, and the breadth of genetic and transgenic mouse models. Overcoming this limitation requires either mouse adaptation of the virus, or heterologous expression of hACE2 in mice. A mouse-adapted strain of SARS-CoV-2 containing 2 amino acid changes in S has recently been described, which results in viral replication in the lungs of mice, and signs of pulmonary damage, but no lethality (26). Alternatively, a number of mouse models for hACE2 expression have been developed including transgenic and knock-in strains, as well as viral vector-mediated delivery of hACE2. Transgenic models include hACE2 expression under the control of the human cytokeratin (K18) epithelial cell promoter (27), the synthetic CAG composite promoter driving high levels of expression in eukaryotic cells (28), human ACE2 promoter (29), or hepatocyte nuclear factor 3/forkhead homologue 4 (HFH4) ciliated epithelial cell promoter (30). A recent report has also described a hACE2 knock-in mouse model (31). Tissue and cellular expression of hACE2 vary in these models leading to differing levels of viral replication in different organs and cell types, and differences in disease pathogenesis. Other models use viral vector-mediated delivery of hACE2 to the lung including adeno-associated virus (AAV) (32) and adenovirus (15, 33). Importantly, a number of these models including K18, CAG, HFH4, and knock-in mice show neuroinvasion and high titer replication in the brain, which likely drives lethality. Although neurological complications have been described in human COVID-19 patients (34), lung damage is the primary cause of death in most patients (35). It is evident that there is an urgent need to establish animal models which authentically recapitulate human lung disease which will be important for understanding disease pathogenesis. However, models in which hACE2 is over-expressed are useful for early vaccine and antiviral studies which use protection from infection or viral replication as an endpoint. Therefore, in this study, we compared viral replication and morbidity in the K18 transgenic hACE2 model head-to-head with adenovirus (Ad)-mediated delivery of hACE2 to the lung.

## Materials and Methods

### Mice

Hemizygous 6-week old female K18-hACE2 mice on the C57BL/6J background (Jax strain 034860), were compared to age and sex-matched wildtype (WT) C57BL/6J (Jax strain 000664) and WT BALB/cJ (Jax strain 000651) mice. Animal studies were approved by the Institutional Animal Care and Use Committee (IACUC) of Icahn School of Medicine at Mount Sinai (ISMMS). Mice were housed in a BSL-2 facility for intranasal instillation of non-replicating adenoviral (Ad) vectors before being transferred to a BSL-3 facility at ISMMS for challenge with SARS-CoV-2. Mice were housed under specific pathogen-free conditions in individually ventilated cages and fed irradiated food and filtered water.

### Cell lines and culture media

T-REx™-293 cells (Life Technologies, Carlsbad, CA) were maintained in high glucose (4500mg/L) Dulbecco Modified Eagle Medium (DMEM) supplemented with 4mM L-glutamine, 100 IU/mL penicillin, 100 μg/mL streptomycin and 10% fetal bovine serum (FBS). Vero-E6 cells (ATCC® CRL-1586™, clone E6) were grown in DMEM containing 10% FBS, non-essential amino acids, 2-[4-(2-hydroxyethyl)piperazin-1-yl]ethanesulfonic acid (HEPES), and penicillin-streptomycin. A549 (ATCC® CCL-185™) cells were cultured in Kaighn’s Modification of Ham’s F-12 (F-12K) containing 10% FBS and penicillin-streptomycin, as above. Cells were maintained at 37°C with 5% CO_2_.

### Viruses

Cells and mice were infected with SARS-CoV-2, isolate USA-WA1/2020 (BEI resources; NR-52281) under BSL-3 containment in accordance to the biosafety protocols developed by the Icahn School of Medicine at Mount Sinai. Viral stocks were grown in Vero-E6 cells in the above media containing 2% FBS for 72 h and were validated by genome sequencing. Cells were infected at a multiplicity of infection (MOI) of 0.1; mice were infected with 1×10^4^ plaque forming units (PFU). Viral seed stocks for non-replicating E1/E3 deleted viral vectors based on human adenovirus type-5 (*HAdV-C5, referred to as Ad throughout*) without an antigen (Ad-Empty), or expressing the human angiotensin-converting enzyme 2 (Ad-ACE2) receptor under the control of a CMV promoter, were obtained from Iowa Viral Vector Core Facility. Viral stocks were amplified to high titers following infection of T-Rex™-293 cells and purification using two sequential rounds of cesium chloride (CsCl) ultracentrifugation, as described previously (36, 37). Infectious titer was determined using a tissue culture infectious dose-50 (TCID_50_) end-point dilution assay, and physical particle titer quantified by micro-bicinchoninic acid (microBCA) protein assay, both described previously (36).

### Flow cytometry and SARS-CoV-2 plaque assay

A549 cells were seeded in 24-well plates at 1×10^5^ cells/well and allowed to adhere overnight. The following day, cells were washed with PBS and transduced with Ad5-Empty or Ad5-hACE2 in triplicate at a MOI of 100 in serum-free Hams F-12K for 3h at 37°C. Control wells were incubated with serum-free Hams F-12K for 3h at 37°C. Following incubation, the suspension was aspirated and media replaced with complete Hams F12-K with 10% FBS. Ad-transduced cells were incubated for 24h before performing flow cytometry staining for surface expression of hACE2 using goat anti-human mAb AF933 (R&D Systems, Minneapolis, MN) used at a final concentration of 10μg/mL. Separate plates, treated identically, were transferred to the BSL-3 facility for subsequent infection with SARS-CoV-2. SARS-CoV-2 plaque assays were performed on cell supernatant or tissue homogenates. 10-fold serial dilutions were prepared in 0.2% BSA/PBS and plated onto a VeroE6 monolayer and incubated with shaking for 1 hr. Inoculum was removed and plates were overlaid with Minimal Essential Media (MEM) containing 2% FBS/0.05% oxiod agar and incubated for 72 hrs at 37°C. Plates were fixed with 4% formaldehyde overnight, stained with a mAb cocktail composed of SARS-CoV-2 spike (Creative-Bios; 2BCE5) and SARS-CoV-2 nucleoprotein (Creative-Biolabs; NP1C7C7) followed by anti-Mouse IgG-HRP (Abcam ab6823) and developed using KPL TrueBlue peroxidase substrate (Seracare; 5510-0030).

### In vivo delivery of virus

For *in vivo* delivery of Ad vectors to the lung, mice were anesthetized by intraperitoneal (*i.p*) injection of ketamine and xylazine diluted in water for injection (WFI; Thermo Fisher Scientific, Waltham, MA). Ad-Empty at 2.5×10^8^ PFU, or Ad-hACE2 at doses of 2.5×10^8^ PFU, 1.0×10^8^ PFU or 7.5×10^7^ PFU, were instilled intranasally (*i.n.*) in a final volume of 50 μL sterile PBS. Untreated, control mice received the same volume of sterile PBS. Mice were transferred to the BSL-3 facility on D3 post-Ad for subsequent challenge with SARS-CoV-2 virus on D5. For SARS-CoV-2 challenge, mice were anesthetized as above and infected with 1×10^4^ PFU in 50 μL of PBS. Mice were sacrificed at day 2 and day 5 post-infection by *i.p.* injection of pentobarbital. 75% of lung and spleen, 50% of brain, and 3 inches of small intestine were homogenized in 1 ml of PBS using ceramic beads. Homogenates were briefly centrifuged and supernatant was immediately used for plaque assays. Separate groups of mice were killed on D5 post-Ad instillation for an assessment of lung histology (*n=3-5 per group*). For lung collection, mice were euthanized by CO_2_ exposure and death confirmed by exsanguination following severing of the femoral artery. After death, the trachea was exposed and lungs inflated with 1.5mL of 10% formalin using a 21G needle fitted to a 3mL syringe. Lungs were removed intact, trimmed carefully and loaded into a tissue embedding cassette. Tissue was fixed overnight in 10% formalin, transferred to PBS after 24h and sent for processing and paraffin embedding at the Biorepository and Pathology Core at ISMMS.

### Histology and hACE2 immunohistochemistry

Paraffin-embedded lung tissue blocks for PBS-treated, Ad-Empty (2.5×10^8^ PFU) or Ad-hACE2 at doses of 2.5×10^8^-7.5×10^7^ PFU, were cut into 5μm sections. Sections were stained with hematoxylin and eosin (H&E) by the Biorepository and Pathology Core, or serial sections (5μm) provided for immunohistochemical (IHC-P) staining for hACE2 as follows; sections were deparaffinized in xylene-free clearing agent, Histo-Clear (Thermo Fisher Scientific, Waltham, MA) and rehydrated using a decreasing ethanol (EtOH) gradient. Endogenous peroxidase activity was blocked by incubating sections for 10min in Bloxall solution (Vector Laboratories, Burlingame, CA) between the first and second 100% EtOH rehydration steps. Antigen retrieval was performed by boiling in sodium citrate (pH 6.0) solution, as described previously (38). Slides were allowed to cool and were washed with PBS prior to blocking of endogenous biotin or avidin binding proteins, using the Avidin/Biotin Blocking Kit (Vector Laboratories, Burlingame, CA) as recommended by the manufacturer. Sections were blocked for 30min at 37°C with blocking buffer provided in the VECTASTAIN® Elite ABC-HRP Kit (Rabbit IgG) with the addition of 1% (w/v) BSA to act as a stabilizer and 0.1% (v/v) Triton X-100 to facilitate permeabilization of the tissue. Following blocking, tissue sections were separated using a hydrophobic pen. A monoclonal rabbit isotype control [clone #SP137; Abcam, Cambridge, MA] or monoclonal rabbit anti-human ACE2 antibody [clone #EPR4436; Abcam Cambridge, MA) diluted in blocking buffer (*see above*) were added to one section on each slide at a final concentration of 1.33μg/mL for 1h at room temperature. Following incubation, sections were washed three times in PBS. Biotinylated anti-rabbit secondary antibody was prepared as instructed by guidelines for the VECTASTAIN® Elite ABC-HRP (Rabbit IgG) kit and sections incubated for 30min at room temperature. Following incubation, sections were washed in PBS and the VECTASTAIN ELITE ABC reagent was prepared and allowed to stand at room temperature for 30min. The pre-made Elite ABC reagent was then added to sections for 30min at room temperature. Cells were washed again three times in PBS and the 3,3’-diaminobenzidine (DAB) chromogen DAB Peroxidase (HRP) Substrate Kit (Vector Laboratories, Burlingame, CA) prepared immediately prior to use and addition to each slide individually. Development of a positive control slide was timed under a microscope and the same development time (10min) applied to all other sections. Sections were transferred to distilled H_2_O and nuclei counterstained for 45s using Haematoxylin QS (Vector Laboratories, Burlingame, CA) before being rinsed extensively with tap water. Sections were then dehydrated by using an increasing gradient of EtOH, ending in Histo-Clear solution. Sections were mounted using Histomount Solution (Life Technologies) and provided to a veterinary pathologist at the Center for Comparative Medicine and Surgery (CCMS) Comparative Pathology Laboratory, ISMMS (*Dr. Virginia Gillespie*). The pathologist evaluated and photographed both IHC-P for hACE2, and H&E sections, and was blinded to the treatment groups. Images were captured under 200X magnification using an Olympus BX43 with the Olympus DP21 Digital Camera system. Representative images from each group are shown alongside a matched isotype control and H&E section.

## Results

### Validation of adenovirus-mediated delivery of hACE2

Recombinant Ad vectors can be used to transduce a wide-range of cell types, both *in vitro* and *in vivo*, and are well-established to facilitate sustained transgene expression when used as a delivery vector (39-41). Prior studies have demonstrated that Ad-mediated delivery of a functional viral entry receptor into mice, such as the human dipeptidyl peptidase 4 (hDPP4) receptor for Middle Eastern Respiratory Syndrome coronavirus (MERS-CoV), can render them permissive to infection with human viruses, which are otherwise unable to support viral entry using the murine receptor homolog (33, 42). In order to validate the functionality of our Ad constructs prior to *in vivo* challenge studies, we transduced A549 cells with an empty adenovirus (Ad-Empty) or an adenovirus expressing hACE2 (Ad-hACE2). We confirmed by flow cytometry staining that transduction of A549 cells with Ad-hACE2 resulted in detectable hACE2 expression on the surface of cells, but no staining was detected following transduction with Ad-Empty (**Fig.1A**). A549 cells do not express detectable levels of ACE2 (43), and it has been reported that they are not permissive to infection with SARS-CoV-1 (43) or SARS-CoV-2 (1). In agreement with published studies, we confirmed that negative control A549, or A549 cells transduced with Ad-Empty failed to support viral replication when infected with SARS-CoV-2 *in vitro*, while A549 cells transduced with Ad-hACE2 supported multicycle viral replication, confirming the functionality of the Ad-hACE2 construct (**Fig.1B**).

**Figure 1.**
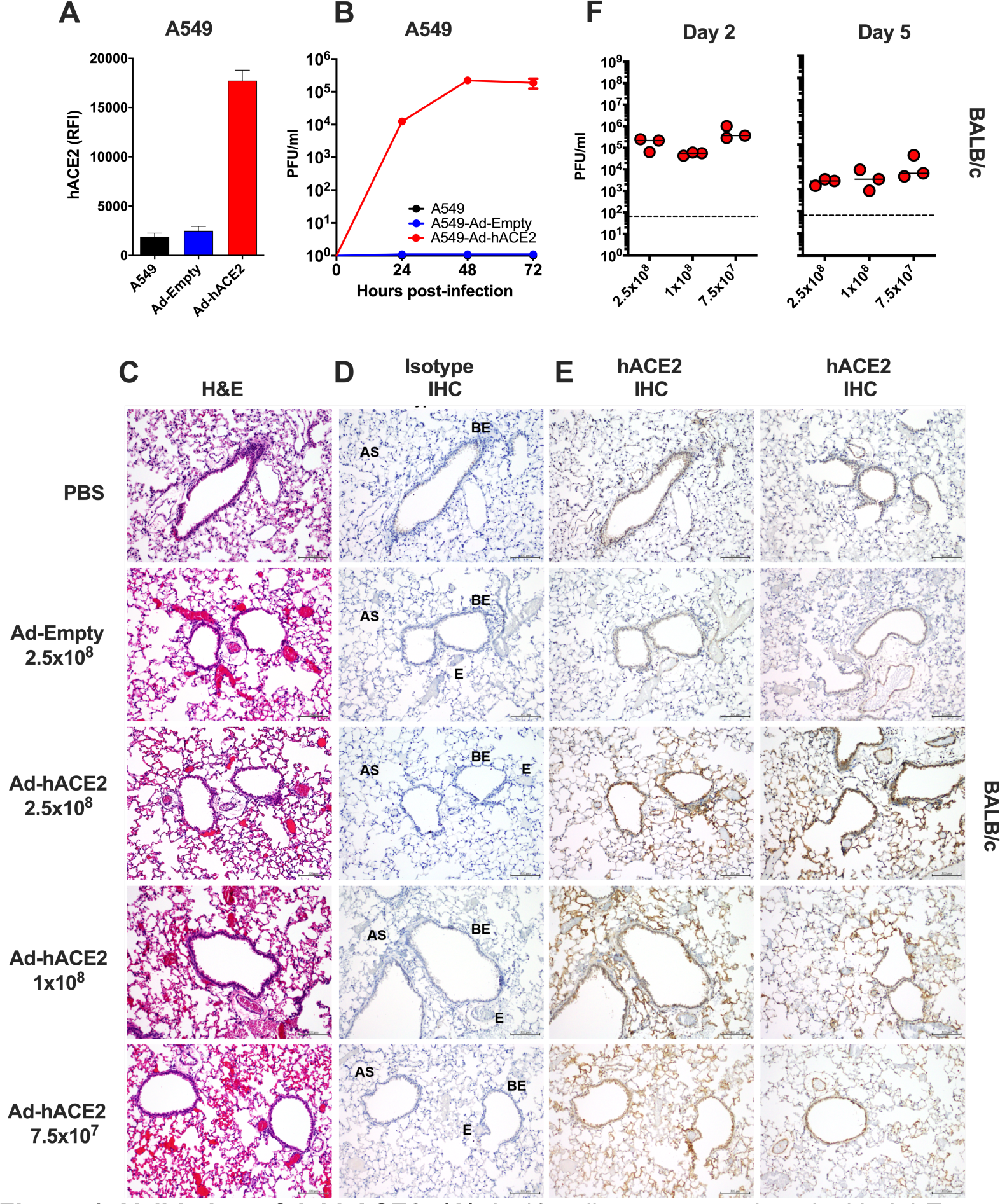
Validation of Ad-hACE2. **(A)** A549 cells were transduced with Ad-Empty or Ad-hACE2 at an MOI of 100 for 3h at 37°C. 24h post-transduction, surface expression of hACE2 was detected by flow cytometry. RFI = relative fluorescence intensity generated by multiplying % hACE2+ cells by the geometric mean fluorescence. **(B)** A549 cells were transduced with Ad vectors as described in (A) followed by infection 24h later with SARS-CoV-2 USA-WA1/2020 at an MOI of 0.1. Virus titers were determined by plaque assay on VeroE6 cells. BALB/c mice were administered intranasally (*i.n*) with the indicated dose of Ad-Empty, Ad-hACE2, or PBS. Lungs were harvested on day 5 post-transduction, paraffin embedded and 5μm sections stained for H&E **(C)**, or for IHC-P using an isotype control **(D)** or α-hACE2 monoclonal antibody **(E)**. Regions of the lung anatomy are indicated on isotype control sections; AS = alveolar septa, BE = bronchiolar epithelium, E = endothelium. Scale bar is 100nm. **(F)** Separate groups of BALB/c mice administered *i.n.* with PBS, Ad-Empty or Ad-hACE2 (2.5×10^8^, 1×10^8^, or 7.5×10^7^ PFU) were infected five days later (D5) with 1×10^4^ pfu of SARS-CoV-2 and lung viral titers were determined by plaque assay on D2 and D5 post-SARS-CoV-2 challenge. *Note;* data points for viral lung titers for BALB/c mice treated with Ad-hACE2 at a dose of 2.5×10^8^ PFU shown in Fig.1F, are the same group of mice as shown in Fig.2D and are shown for visualization purposes to allow a comparison between doses, although all groups were part of the same larger experiment.

To over-express hACE2 in the lungs of mice, we next administered BALB/c mice intranasally (*i.n*) with PBS, 2.5×10^8^ PFU of Ad-Empty, 2.5×10^8^, 1×10^8^ or 7.5×10^7^ PFU of Ad-hACE2. Innate immune responses to significantly higher doses of Ad vectors (1×10^11^) are known to peak between 6-24h post-administration (36, 44). Therefore, we reasoned that *in vivo* challenge with SARS-CoV-2 at D5 post-Ad transduction would be minimally impacted by non-specific inflammatory responses. In support of this, the D5 time-point has also recently been used successfully for Ad-mediated delivery of viral receptors to the lungs of mice by other investigators (33, 42). Five days post-transduction (D5), lungs were harvested for an evaluation of histology by H&E staining, and by immunohistochemistry for hACE2 expression. All sections were assessed and evaluated by a veterinary pathologist who was blinded to the treatment groups. H&E staining did not reveal overt signs of inflammation due to administration of the Ad vectors and in general all sections were considered to be very similar (**Fig.2C**). Specimens from all treatment groups were reported to display diffuse, alveolar congestion with multifocal, acute hemorrhage comprising <25% of the section, along with scant to mild fibrin deposition which was largely localized to the alveolar septa. These effects were considered to be potentially associated with euthanasia using inhaled CO_2_ (45). One H&E section in the 7.5×10^7^ PFU Ad-hACE2 group and one in the 2.5×10^8^ PFU Ad-hACE2 group exhibited multifocal lymphoid aggregates and a few scattered perivascular and peribronchiolar lymphocytes and plasma cells. The latter section also had evidence of focal bronchus-associated lymphoid tissue (BALT). Random lymphoid aggregates with no other associated lesions in multiple tissues including the lungs are common background observations in mice (45). Therefore, any inflammation induced by the Ad vectors at the doses tested was considered to be mild.

**Figure 2.**
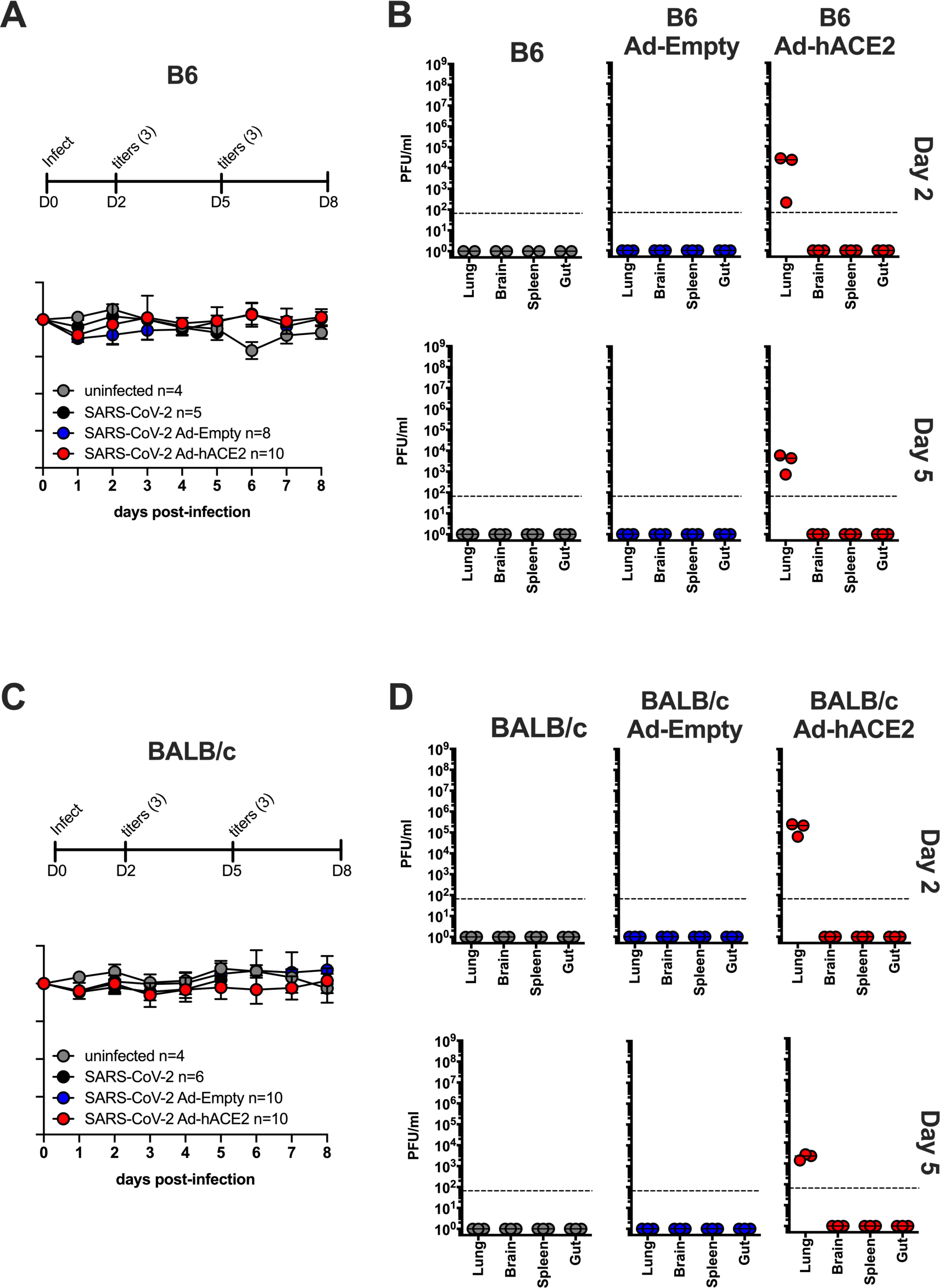
B6 and BALB/c Ad-hACE2 SARS-CoV-2 infection. **(A-B)** B6 mice and **(C-D)** Balb/c mice were transduced with 2.5×10^8^ PFU of Ad-empty, Ad-hACE2, or PBS. On D5 post-Ad administration mice were infected with 1×10^4^ pfu of SARS-CoV-2 and monitored for weight loss (A and C) and viral titers (B and D) according to the indicated timeline. n=animal number at day 0, as animals were harvested for titers n was reduced according to the diagram. *Note;* data points for viral lung titers for Balb/c mice treated with Ad-hACE2 at a dose of 2.5×10^8^ PFU shown in Fig.2D, are the same group of mice as shown in Fig.1F.

Sections from all treatment groups (PBS control, Ad-Empty at 2.5×10^8^ PFU and Ad-hACE2 at increasing doses) were stained using anti-hACE2 or species-matched isotype control mAb simultaneously (**Fig.2D, E**). The anti-hACE2 monoclonal antibody (mAb) we used for IHC-P detection of hACE2 is specific for hACE2 and lacks cross-reactivity to mouse tissue or to murine ACE2. No staining was observed for the isotype control antibody (**Fig.2D**). PBS control sections were determined by the pathologist to be negative for hACE2 staining. All Ad-hACE2 treated mice displayed a similar profile of hACE2 staining in the lung with rare to segmental labeling of bronchiolar epithelial (BE) cells, multifocal patchy labeling of alveolar septa (AS) and labeling of few alveolar macrophages (*data not shown*). In some sections, staining was also observed on the endothelium and/or walls of few medium to small blood vessels. Positive staining was patchy, as would be expected for non-uniform transduction of the lung with the Ad-hACE2 vector. Transduction of epithelial cells, alveolar macrophages and endothelial cells is consistent with the known tropism and target cells for HAdV-C5 based vectors in mice (36, 39, 46, 47). Although the lowest dose of Ad-hACE2 (7.5×10^7^ PFU) appeared to yield a similar distribution of transduction in the lung to the highest dose (2.5×10^8^ PFU), the intensity of staining appeared to be dose-dependent with more intense brown staining for hACE2 in the highest dose of Ad-hACE2 (2.5×10^8^ PFU).

In order to determine the optimal Ad-ACE2 dose for SARS-CoV-2 replication *in vivo*, separate groups of BALB/c mice were administered with 2.5×10^8^ (*data also shown in Fig.2D*), 1×10^8^, or 7.5×10^7^ PFU of Ad-ACE2 followed by infection with SARS-CoV-2 at D5 post-Ad transduction. Viral lung titers were similar among the different doses, with slightly higher replication at the lowest Ad-ACE2 dose (**Fig.1F**), indicating that doses as low as 7.5×10^7^ PFU of adenovirus are sufficient to confer hACE2 expression and support SARS-CoV-2 replication.

### B6 and BALB/c Ad-hACE2 SARS-CoV-2 infection

In parallel, we used our adenovirus constructs to transduce both B6 and BALB/c mice intranasally with 2.5×10^8^ PFU of Ad-hACE2 or Ad-Empty. At 5-days post-transduction mice were infected with 1×10^4^ pfu of SARS-CoV-2 and monitored for body weight loss. Significant weight loss was not observed in any of the groups on the B6 (**Fig.2A**) or Balb/c (**Fig.2C**) backgrounds at this dose of SARS-CoV-2, although mild weight loss (<10%) was observed in an unrelated experiment when mice were challenged with a 5-fold higher dose of SARS-CoV-2 (*data not shown*). At day 2 and day 5 post-infection, organs were harvested from mice to determine viral titers. Ad-hACE2 transduced mice on both the B6 and BALB/c background showed viral titers in the lung at day 2 post-infection, with titers decreasing at day 5. Viral titers in BALB/c mice were ∼1 log higher at day 2 compared to B6 mice (**Fig.2B** and **2D**). No viral replication was observed in the brain, spleen, or gut, consistent with *i.n.* administration of a relatively low dose of a HAdV-C5-based Ad vector and established knowledge regarding the lack of significant tropism for these organs in mice (48-50). WT B6, BALB/c, B6 Ad-Empty, and BALB/c Ad-Empty mice failed to support SARS-CoV-2 replication as expected based on lack of hACE2 receptor expression.

### K18-hACE2 SARS-CoV-2 infection

B6 K18-hACE2 mice were similarly infected with 1×10^4^ pfu of SARS-CoV-2 and monitored for weight loss and viral titers. K18-hACE2 mice showed progressive weight loss – by day 7 one animal was below 75% body weight and by day 8 one animal was found dead (**Fig.3A**). K18-hACE2 mice showed high viral titers in the lung at day 2 post-infection, and some animals had virus in the brain. By day 5 post-infection viral titers in the lung were reduced while titers in the brain had increased. Additionally, virus was detectable in the gut by day 5 (**Fig.3B**). Notably, viral titers in the lung were 2-3 logs higher compared to Ad-hACE2 SARS-CoV-2 infected mice. Replication of SARS-CoV-2 to high titers in multiple organs likely reflects the expression of hACE2 in cells of these organs in this transgenic mouse model (27).

**Figure 3.**
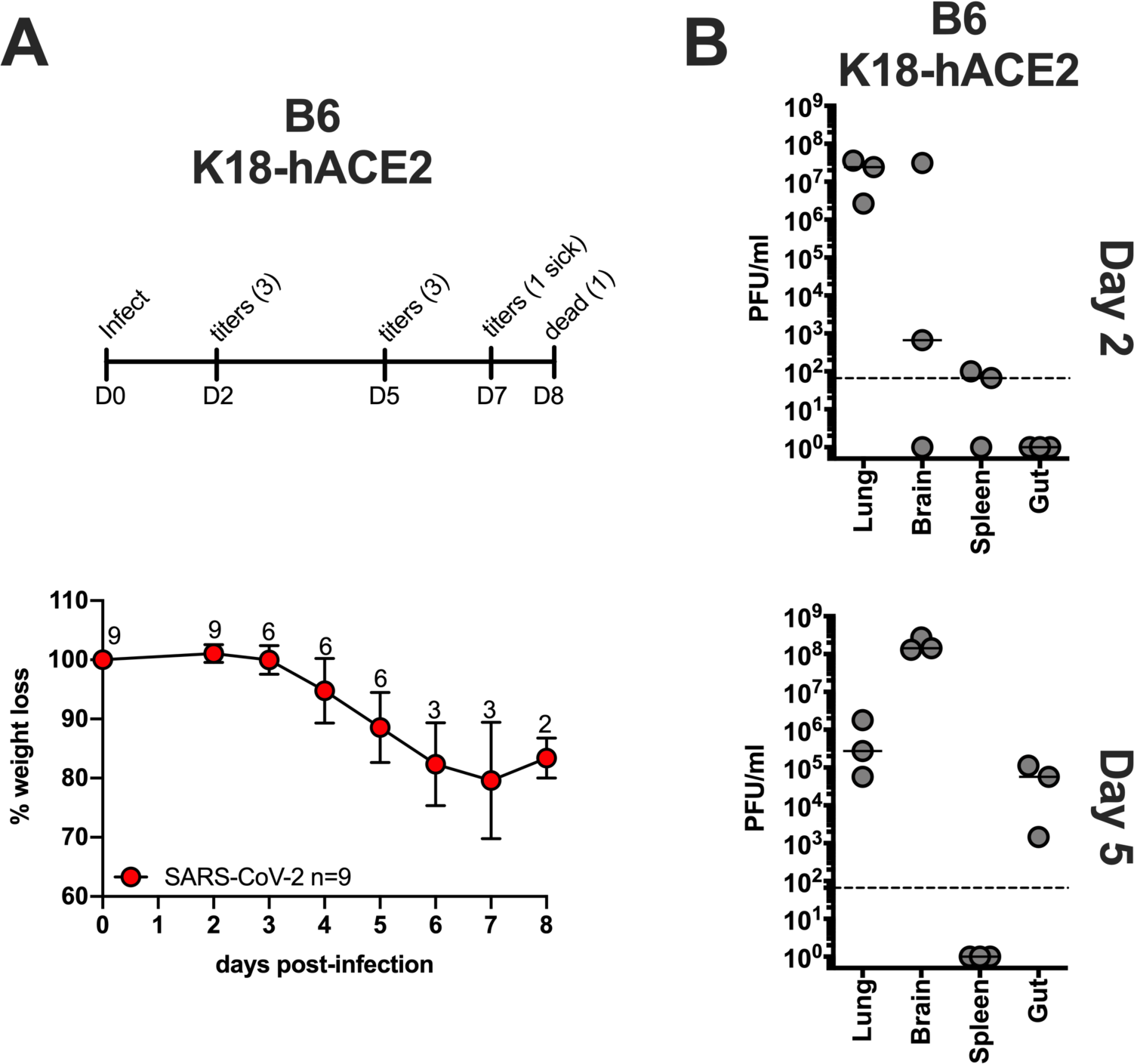
B6-K18-hACE2 SARS-CoV-2 infection. B6 K18-hACE2 mice were infected with 1×10^4^ PFU of SARS-CoV-2 and monitored for weight loss **(A)** and viral titers **(B)** according to the indicated timeline. n=animal number at day 0, as animals were harvested for titers n was reduced according to the diagram.

## Discussion

The development of small animal models which are permissive to infection with SARS-CoV-2 will be invaluable, and will not only allow us to better understand the pathogenesis of the virus, but will enable characterization of immune responses to vaccine platforms or immunization regimens which confer protection. The rapid emergence of the SARS-CoV-2 virus prompted the research community to try to apply information gained from historical studies of SARS-CoV-1 and other coronaviruses to the development of a relevant animal model for SARS-CoV-2. It was already established in the literature that Syrian hamsters were permissive to infection with SARS-CoV-1 (51-53), therefore it was reasonable that they might also represent a useful model for SARS-CoV-2 infection. Indeed, Syrian hamsters have recently been reported to authentically model clinical features of severe human COVID-19 pathogenesis, including alveolar damage and bilateral interstitial pneumonia (54). In addition, hamsters mounted neutralizing antibody responses to SARS-CoV-2 which were capable of providing protection from subsequent virus re-challenge. However, phenotypic or functional studies of T cell responses or other immune cell populations to immunization or infection in Syrian hamsters may be limited by the lack of availability of reagents, particularly when compared with mice. Therefore, the development of a permissive mouse model would facilitate broad and adaptable applications to evaluate pathogenesis and replication, and uncover the underlying factors that contribute to immunopathology. Mouse models offer a number of benefits in this context, which include the accessibility of inbred genetic backgrounds on multiple strains, large numbers of genetically-modified strains, existing models of co-morbidities such as obesity or diabetes (55), and commercial access to many antibodies and tools for immunological studies.

Prior studies to validate a mouse model for other coronaviruses had reported the use of transgenic models to express coronavirus receptors (27, 42, 56-60). Achieving this for MERS-CoV has parallels with SARS-CoV-2 in the sense that standard laboratory strains of mice are not permissive to infection, as a result of their inability to use the murine ortholog of their entry receptors, hDPP4 and hACE2, respectively. Studies in which the homologs of these receptors were delivered either to non-permissive cells *in vitro* (43), or *in vivo* to the mouse lung by overexpression via viral vectors (32, 42), could sensitize to infection. Since the emergence of the SARS-CoV-2 virus, investigators have rapidly responded to the pandemic by evaluating the potential for already-generated (58), but not widely available, hACE2 transgenic mice in supporting SARS-CoV-2 infection and pathogenesis (29, 30). Although useful, these models produced variable results in which some supported low level SARS-CoV-2 replication with minimal morbidity or lung pathology (29), and others exhibited signs of local and systemic disease which partially mimic symptoms of severe human COVID-19 infection, including increased mortality in male mice (30). Several recent reports have also now described the success of the Ad-hACE2 vector delivery approach in mice, and despite minor differences in the approach, our results strongly support their findings (33, 61). We, and others, detected hACE2 expression predominantly on alveolar epithelial cells, with some expression in bronchiolar epithelial cells following transduction with a dose of 2.5×10^8^ PFU Ad-hACE2 (33, 61). In this study, we also detected some transduction in endothelial cells and alveolar macrophages, which is consistent with the known *in vivo* cellular tropism for Ad vectors based on HAdV-C5 in mice (36, 39, 49, 50).

The K18-hACE2 transgenic model has been well characterized for SARS-CoV-1 (27, 43), but to our knowledge has not yet been comprehensively evaluated for SARS-CoV-2 infection, in terms of measuring morbidity and viral dissemination with subsequent replication in extra-pulmonary organs. Therefore, in this study, we wanted to perform a head-to-head comparison of the K18-hACE2 model and the Ad-hACE2 murine model in supporting SARS-CoV-2 viral replication in both BALB/c and B6 mice in parallel. We also use different amounts of the Ad-hACE2 vector to determine if titers below 2.5×10^8^ PFU could be used to support SARS-CoV-2 challenge. We reasoned that a comparison of these different hACE2 model systems would provide useful information for the research community in deciding which model to apply to specific research hypotheses. Our findings have determined that while both the K18-hACE2 and Ad-hACE2 models support SARS-CoV-2 replication, they have a number of advantages and disadvantages. SARS-CoV-2 replication in the lungs of K18-hACE2 mice following a challenge dose of 1×10^4^ PFU was substantially higher than B6 mice (same genetic background) and BALB/c mice administered with Ad-hACE2 vectors. In addition, the K18-hACE2 model resulted in more severe disease, manifesting in weight loss, and replication in multiple organs – including lung, brain, and gut. Therefore, the K18-hACE2 mice could provide a stringent model for testing vaccines and antivirals, where weight loss, mortality or extra-pulmonary viral replication are end-points. However, brain infection and neurological complications, while reported in humans, are not the primary cause of death in most human COVID-19 patients (35). While we did not observe weight loss in the Ad-hACE2 model, a recent study using the same model did report weight loss using a higher infectious dose (1×10^5^ FFU compared to 1×10^4^ PFU in our study). Other possible reasons for discrepancies could include differences in routes of inoculation of SARS-CoV-2 (both *i.n.* and *i.v.* compared to only *i.n.* in our study) (33). However, we have recently generated data in a separate (unpublished) study which demonstrates mild weight loss when a challenge dose of 5×10^4^ PFU is used.

Some studies using Ad-hACE2 have used IFNAR^-/-^ mice, or have blocked IFNAR1 using α-IFNAR1 mAbs, demonstrating that a lack of intact type I IFN response in mice resulted in greater weight loss and exacerbated lung pathology upon SARS-CoV-2 infection (15, 33), and concluding that IFN may have protective effects against SARS-CoV-2. However, this should be validated in an additional model for hACE2, as there is evidence that blockade of Type I IFN can affect transgene expression from non-replicating Ad vectors and T cell responses to the transgene/vector, which could confound results (39, 62). Other caveats of the Ad-hACE2 animal model include the non-uniform transduction of the lung epithelium (and consequently, non-uniform expression of hACE2 in the mouse lung), the possibility of triggering non-specific inflammatory responses if used at doses higher than 2.5×10^8^ PFU – or with poorly prepared Ad stocks which have large quantities of empty capsids, and the potential lack of suitability for use of the Ad-hACE2 model in immunization studies using non-replicating Ad vaccines for SARS-CoV-2. However, with this in mind, we describe the successful use of a dose of Ad-hACE2 (7.5×10^7^ PFU) which supports equivalent replication of SARS-CoV-2 in the lung to the previously reported higher dose (33, 61). Most importantly, the flexibility of the Ad-hACE2 model allows studies of multiple mouse strains immediately, without time-consuming breeding to a hACE2 transgenic or knock-in background. In conclusion, further refinement and development of these models and additional small animal models will be critical for studying disease pathogenesis, and for evaluating novel therapeutics and vaccines to protect against SARS-CoV-2 infection.

## Acknowledgements

This work was partially supported by the National Institute of Allergy and Infectious Diseases (NIAID) Centers of Excellence for Influenza Research and Surveillance (CEIRS) contract HHSN272201400008C (F.K., A.G.-S.), by supplements to NIAID grant U19AI135972 and DoD grant W81XWH-20-1-0270 (A.G.-S.), by the generous support of the JPB Foundation (A.G.-S.), the Open Philanthropy Project (research grant 2020-215611 (5384)) and other philanthropic donations to A.G.-S and F.K, and by NIAID R21AI157606 (L.C). Viral Vectors were provided by the University of Iowa Viral Vector Core, and we thank Susan Stamnes, Kaylee Murphy (Iowa Viral Vector Core) and Dr. Paul B. McCray Jnr (University of Iowa) for making the Ad5-hACE2 virus rapidly available to us. We thank Alan Soto and Frances Avila at the Biorepository and Pathology Core (ISMMS) for tissue processing and histology. We thank staff at the Center for Comparative Medicine and Surgery (CCMS) Icahn vivarium, Carlos Franco, Lenny Martinez and Joseph Espinoza, for their assistance in coordinating transfer of animals to the BSL-3 facility. We also thank Randy Albrecht for support with the BSL3 facility and procedures at the ISMMS.

